# Large genomic variants reveal unexplored intraspecific diversity in *Brassica rapa* genomes

**DOI:** 10.1101/2020.07.02.183640

**Authors:** Julien Boutte, Loeiz Maillet, Thomas Chaussepied, Sébastien Letort, Jean-Marc Aury, Caroline Belser, Franz Boideau, Anael Brunet, Olivier Coriton, Gwenaëlle Deniot, Cyril Falentin, Virginie Huteau, Maryse Lodé, Jérôme Morice, Gwenn Trotoux, Anne-Marie Chèvre, Mathieu Rousseau-Gueutin, Julie Ferreira de Carvalho

**Affiliations:** IGEPP, INRAE, Institut Agro, Univ Rennes, 35653 Le Rheu, France; IRISA/INRIA, Campus de Beaulieu, 35042 Rennes Cedex, France; Génomique Métabolique, Genoscope, Institut de biologie François-Jacob, CEA, CNRS, Univ Evry, Université Paris-Saclay, 91057 Evry, France

**Keywords:** *Brassica*, Genome evolution, Transposable elements, LTR Gypsy, ribosomal DNA, Intraspecific diversity

## Abstract

Traditionally, reference genomes in crop species rely on the assembly of one accession, thus occulting most of intraspecific diversity. However, rearrangements, gene duplications and transposable element content may have a large impact on the genomic structure, which could generate new phenotypic traits. Using two *Brassica rapa* genomes recently sequenced and assembled using long-read technology and optical mapping, we investigated structural variants and repetitive content between the two accessions and genome size variation among a core collection.

We explored the structural consequences of the presence of large repeated sequences in *B. rapa* ‘Z1’ genome versus the *B. rapa* ‘Chiifu’ genome, using comparative genomics and cytogenetic approaches. First, we showed that large genomic variants on chromosomes A05, A06, A09 and A10 are due to large insertions and inversions when comparing *B. rapa* ‘Z1’ and *B. rapa* ‘Chiifu’ at the origin of important length differences in some chromosomes. For instance, lengths of ‘Z1’ and ‘Chiifu’ A06 chromosomes were estimated *in silico* to be 55Mb and 29Mb, respectively. To validate these observations, we compared using fluorescent *in-situ* hybridization (FISH) the two A06 chromosomes present in a F1 hybrid produced by crossing these two varieties. We confirmed a length difference of 17.6% between the A06 chromosomes of ‘Z1’ compared to ‘Chiifu’. Alternatively, using a Copy Number Variation approach, we were able to quantify the presence of a higher number of rDNA and Gypsy elements in ‘Z1’ genome compared to ‘Chiifu’ on different chromosomes including A06. Using flow cytometry, the total genome size of 12 *Brassica* accessions corresponding to a *B. rapa* available core collection was estimated and revealed a genome size variation of up to 16% between these accessions as well as some shared inversions.

This study revealed the contribution of long-read sequencing of new accessions belonging to different cultigroups of *B. rapa* and highlighted the potential impact of differential insertion of repeat elements and inversions of large genomic regions in genome size intraspecific variability.

## INTRODUCTION

Genetic and phenotypic diversity are the drivers of plant evolution and adaptation. While ecology and population genetics provide important knowledge on the molecular processes underlying the astonishing plant biodiversity, comparative genomic analyses are essential to understand the large-scale genomic variations that can be observed within species and how these structural variations may impact plant phenotypes. Re-sequencing of accessions and construction of pangenomes of plant species such as wheat (Montenegro et al., 2017), maize (Gage et al., 2019), rice (Schatz et al., 2014; Wang et al., 2018; Zhao et al., 2018) or soybean (Lam et al., 2010; Li et al., 2014) have revealed the involvement of structural variations in various agronomically relevant traits. In maize, structural variants are implicated in plant architecture, flowering and disease resistance (Walker et al., 1995; Chia et al., 2012; Lu et al., 2015). Similarly, using pangenomes, numerous structural variations have been revealed in *Brassica napus* varieties correlating with genes and phenotypic traits such as silique length and seed weight (Gazave et al., 2016), disease resistance (Dolatabadian et al., 2020) as well as flowering time and vernalization (Song et al., 2020). Structural variants identified as underlying flowering time were associated with the insertion of transposable elements in three FLC genes (Song et al., 2020). However, structural diversity in the progenitors of this allopolyploid species have so far been particularly overlooked.

Only recently, the development of *Brassica* pangenomes has started to reveal the extensive number of structural variants within one of the two diploid progenitors: *Brassica oleracea* (Golicz et al., 2016) and assessed in regards to the domestication process of the subsequent allotetraploids *B. juncea* (Paritosh et al., 2019) and *B. napus* (Song et al., 2020). These studies highlight the importance of structural variants and gene copy number variation in adaptive phenotypes. While core genes represent 80% of the assembled genomes, each new accession brings around 20% of novel genes. In most of these analyses however, the occurrence and importance of the repeat compartment is undervalued.

Transposable elements form an important internal source of genetic diversity as a result of their ability to create mutations, alter gene expression, and promote chromosomal mispairing (Kidwell and Lisch, 2000). These ancient, ubiquitous, and dynamic components of eukaryotic genomes comprise up to 58% of the allopolyploid genome of *B. napus* (Song et al., 2020). Transposable elements comprise two classes that have contrasted mode of replication, each divided in several families: Retrotransposons (Class I) transpose via reverse transcription of their messenger RNA and DNA Transposons (Class II) encompassing all other types of TEs (Wicker et al., 2007). Overall, TEs have major impact on genome organization and function, and have been associated with up-regulated genes in response to various abiotic stresses, environmental gradients, reproductive mode (asexuality, selfing vs. outcrossing), interspecific hybrid formation and polyploidy (Wright and Schoen, 1999; Kalendar et al., 2000; Ungerer et al., 2006; Makarevitch et al., 2015; Vicient and Casacuberta, 2017). Several studies from different biological systems indicate that the patterns (position and number of copies) of a single Transposable Element (TE) family can vary within species (Quadrana et al., 2016). However, the study of TEs inserted in complex genomes of various populations is challenging and rely on the amplification of specific TE sequences (for instance using Sequence Specific Amplification Polymorphism transposon display, Parisod et al., 2009) or low-coverage sequencing strategies (Ferreira de Carvalho et al., 2016). The utilization of long-read sequencing and optical mapping in several varieties of the same species is allowing fine comparative genomic analysis of the differential proliferation of TEs and other repeats. This overlooked component of plant genomes is getting accessible and provides important insights on the structural genomic diversity standing within species.

In the present study, we conducted a fine comparative genomic analysis of two varieties of *Brassica rapa* ‘Chiifu’ and ‘Z1’. *Brassica rapa* (AA, 2n□=□20) is one of the diploid progenitors of the important allotetraploid oilseed crops, *B. juncea* (AABB, 2n□=□36) and *B. napus* (AACC, 2n□=□38) (U, 1935). Therefore, *B. rapa* varieties can be used to broaden the genetic diversity of these domesticated species. *Brassica* diploid species arose from a Whole Genome Triplication (WGT) that occurred about 15-20 million years ago (Murat et al., 2015). Diploid *Brassica* are thus considered as paleohexaploids. Thereafter, their duplicated genomes rapidly underwent extensive gene deletion through a fractionation process (Murat et al., 2015). The majority of the genes are tending towards a return to single-copy status (Wang et al., 2011), and overall gene content in *Brassica* diploid genomes has so far been reduced by about half. Yet, different cultigroups within *Brassica* species have selected different alleles from the same duplicated genes explaining part of the astonishing and extensive number of morphotypes existing within each species (Cheng et al., 2014, 2016b; Golicz et al., 2016). *Brassica rapa* speciation started after the divergence with *B. oleracea* estimated 2.18 Mya (Li et al., 2017) but its diversification occurred mainly during the domestication process of *B. rapa* crops, 3500 years BC in the Mediterranean basin with a secondary center of diversification in Asia (Guo et al., 2014). This resulted in genetically (Aissiou et al., 2018) and phenotypically diverse morphotypes (leafy, root or fodder and oilseed types) corresponding to different subspecies including ssp. pekinensis (Chinese cabbage with ‘Chiifu’) and ssp. trilocularis (Yellow sarson with ‘Z1’). Chinese cabbage has originated in central China by natural hybridization between Pak-choi (ssp. chinensis) and turnip rape (ssp. oleifera) 1,200-2,100 years ago (Song et al., 1990; Qi et al., 2017). Yellow sarson originated from India but its precise origin is still unknown. Still, the underlying structural genomic diversity has never been explored within this important crop.

Using established protocols (flow cytometry and cytogenetics) along with state-of-the-art whole genome sequencing and optical mapping for assembly of complex *Brassica* genomes, we compared structural rearrangements between two varieties of *B. rapa* that have contrasted morphotypes, evolutionary history and reproductive mode. Specifically, we identified differential insertion of ribosomal DNA and LTR retrotransposons and reconstructed the evolution of these repeats during *B. rapa* domestication. Finally, we unraveled wide variation in DNA content among different accessions belonging to *B. rapa* core collection.

## MATERIALS AND METHODS

### Biological material

Two *B. rapa* genotypes, *B. rapa* var. *pekinensis* cabbage ‘Chiifu’ (pure line) and *B. rapa* var. *trilocularis* ‘Z1’ (doubled haploid line) as well as their F1 hybrid (‘Z1‘x‘Chiifu’) were grown in semi-controlled conditions in a greenhouse. Seven to ten individuals of each *B. rapa* accession representing biological replicates were harvested for leaves and roots. This material will further be used in flow cytometry and fluorescent *in-situ* hybridization (FISH) experiments, respectively. In addition, ten accessions representing the different sub-species of *B. rapa* (Zhao et al., 2010; Cheng et al., 2016b) were grown in a similar environment for assessing genome size variability using flow cytometry. Fresh leaves from three to eight individuals, used as biological replicates, were harvested per accession: from group 1, one accession of *ssp. pekinensis*, from group 2, one accession of *ssp. narinosa;* from group 3, one accession of *ssp. perviridis* and one accession of *ssp. nipposinica;* from group 4, two different accessions of *ssp. oleifera;* from group 5, one accession of *ssp. trilocularis;* from group 6, three different accessions of *ssp. rapa*. Leaf tissue is then directly processed for flow cytometry analyses (Supplementary Table 1).

### Annotations of Transposable Elements

The latest genome assemblies available for *B. rapa* ‘Z1’ (Belser et al., 2018) and ‘Chiifu’ V3 (Zhang et al., 2018) were retrieved from the Genoscope website (http://www.genoscope.cns.fr/plants) and the *Brassica* database (http://brassicadb.org/) respectively. These genomes were subjected to the same transposable element annotation package REPET V2.5 (Quesneville et al., 2005; Flutre et al., 2011; Hoede et al., 2014). Briefly, the REPET pipeline was used for the detection, classification (TEdenovo) and annotation of TEs (TEannot). Using each reference genome, the TEdenovo pipeline identifies TE copies and provides a consensus sequence when at least five copies of the same TE family are retrieved within the genome. Then, TEannot pipeline uses the consensus sequences to annotate the provided reference genomes. Two files containing the structural annotation of ‘Z1’ and ‘Chiifu’ were output in GFF3 format (Generic feature Format version 3). The resulting GFF3 files were hand curated using custom python scripts (Python version 2.7.12).

### Comparative structural analyses

To evaluate large-scale genome structural rearrangements, the assembled genomes of *B. rapa* ‘Z1’ and ‘Chiifu’, as well as the A subgenome of *Brassica napus* Darmor Bzh V4.1 were retrieved from the *Brassica* database (http://brassicadb.org/) and compared. The package MUMmer V3.23 (Kurtz et al., 2004) was used on each of the 10 chromosomes using the following settings: -id 95, -l 4000. Inversions or deletions observed in ‘Z1’ when compared to ‘Chiifu’ were further verified using BioNano raw data from Belser et al., (2018). To better characterize these genomic regions, gene annotations were retrieved from GFF files. GO enrichment was performed on GO terms of the identified genes present in specifically inserted regions of ‘Z1’ using Agrigo V2 (Tian et al., 2017). Finally, we assessed the duplication status of these genes using PCK (Murat et al., 2015) and custom made python scripts. In addition, among all inversions three showing defined breakpoints in both ‘Z1’ and ‘Chiifu’ were chosen and validated using custom-made PCR primers designed in flanking regions of inversions and specific to either ‘Z1’ or ‘Chiifu’ accessions (Supplementary Table 2).

### Validation of rearrangements using cytogenetics

To validate the presence of repeated sequences on the chromosome A06 of *B. rapa* ‘Z1’, fluorescent *in-situ* hybridization (FISH) was carried out in ‘Z1’, ‘Chiifu’ and the F1 hybrid ‘Z1’x’Chiifu’ according to protocols detailed in (Książczyk et al., 2011). The two BAC clones KBrB022P06 and KBrH003P24 hybridizing on each of the A06 chromosome arms in *B. rapa* (Xiong and Pires, 2011) were labelled by random priming with biotin-14-dUTP (Invitrogen, Life Technologies). The ribosomal probe used in this study was pTa-71 (Gerlach and Bedbrook, 1979) which contained a 9-kb *Eco*RI fragment of rDNA repeat unit (18S-5.8S-26S genes and spacers) isolated from *Triticum aestivum*. pTa-71 was labelled with Alexa-488 dUTP by random priming. Biotinylated probes were immunodetected by Texas Red avidin DCS (Vector Laboratories) and the signal was amplified with biotinylated anti-avidin D (Vector Laboratories). The chromosomes were mounted and counterstained in Vectashield (Vector Laboratories) containing 2.5μg/mL 4’,6-diamidino-2-phenylindole (DAPI). Fluorescence images were captured using a ORCA-Flash4 (Hamamatsu, Japan) on an Axioplan 2 microscope (Zeiss, Oberkochen, Germany) and analyzed using Zen 2 PRO software (Carl Zeiss, Germany). Finally, the lengths of both A06 chromosomes in the hybrid ‘Z1’x’Chiifu’ were estimated on 14 cells at the mitotic metaphase stage and measured using the visualization software Zen 2 PRO (Carl Zeiss, Germany).

### Copy number estimation of highly repeated sequences: ribosomal DNA and Gypsy LTR retrotransposon

Copy Number Variations (CNV) of 45S rDNA (otherwise called 25S or 35S rDNA) and Gypsy LTR Retrotransposons were obtained using Shotgun data for *B. rapa* var. pekinensis ‘Chiifu’ (from http://brassicadb.org) and *B. rapa* var. trilocularis ‘Z1’ (from http://www.genoscope.cns.fr/externe/plants/datasets.html), applying a similar method as described in Boutte et al. (in prep) based on a calibration using the CNV (based on read depth approach) of single copy genes. First, 17,954 putative single copy genes in ‘Chiifu’ and ‘Z1’ were identified using custom made python scripts based on PCK genes from *B. rapa* (Murat et al., 2015). Second, 38 rDNA sequences, corresponding to *Brassica* rDNA full length sequences downloaded from the NCBI (last accessed August 5th, 2019) and 96 Gypsy sequences downloaded from the Gypsy database (http://gydb.org, last accessed August 5th, 2019) were retrieved as reference sequences. Then, rDNA and Gypsy sequences of ‘Chiifu’ and ‘Z1’ were identified by performing BLASTn and tBLASTn algorithms (*e*-value threshold of 10^−5^; Altschul et al., 1997) between the rDNA and Gypsy reference sequences and the two assembled genomes ‘Chiifu’ v.3.0 (Zhang et al., 2018) and ‘Z1’ (Belser et al., 2018). Finally, consensus sequences (two rDNA and 211 Gypsy) were obtained overall for ‘Z1’ and ‘Chiifu’ using python custom script (sequences presenting more than 90% identity were combined together) and aligned with Mafft (v.7.245; --auto option, (Katoh and Standley, 2013). Visual checking of the alignments was done using Geneious v.10.0.9 (Kearse et al., 2012). ‘Chiifu’ and ‘Z1’ genomic reads were then mapped on the two 45S rDNA consensus, the 211 Gypsy RT consensus and the 17,954 putative single copy genes using Bowtie 2, v.2.2.7 (Langmead and Salzberg, 2012) using the parameters G, 52, 6 and G, 58, 3 for ‘Chiifu’ and ‘Z1’, respectively. CNV of 45S rDNA and Gypsy sequences were estimated using mapping results and previously developed custom python scripts (Boutte et al. in prep).

### Phylogenetic analyses

To study the evolutionary relationships of 45S rDNA and Gypsy sequences of *Brassica rapa* ‘Chiifu’ and ‘Z1’, phylogenetic analyses were performed. First, 45S rDNA and Gypsy sequences (firstly translated to protein sequences and merged to the 96 Gypsy database sequences) of ‘Chiifu’ and ‘Z1’ were aligned separately using Mafft alignment (v.7.245; --auto option, (Katoh and Standley, 2013). Second, beginning and end of the matrices with less than 25% of the sequences were deleted as well as indels present in more than 25% of the sequences. Phylogenetic analyses were performed using Geneious tree builder with the Jukes-Cantor model and the Neighbour-joining method (45S rDNA clade support was determined by 100 bootstrap replicates). To avoid Long Branch Attraction error, sequences presenting long branches were deleted and new phylogenetic analyses were performed using the parameters described above.

### Genome size estimation by flow cytometry

In order to explore the intraspecific variability of genome size among the different varieties of *B. rapa*, we assessed by flow cytometry, the DNA content and estimated genome size for the focal accessions ‘Chiifu’ and ‘Z1’ as well as ten other accessions representing *B. rapa* intraspecific diversity. Briefly, approximately 4mg of fresh leaves of *B. rapa* and *Pisum sativum* (var. Victor; used as an internal reference standard) were harvested and transferred to a Petri dish. This material was chopped using a sharp razor blade in 500μl of staining buffer (from Cystain PI OxProtect) and incubated at room temperature for 30sec to 90sec. The solution was then filtered through a 50μm nylon mesh and 1.5ml of solution (0.0166mg of RNase A and 10μl of Propidium Iodide) was added per sample. Incubation at room temperature was made for 30min to 60min, protected from light. Estimation of genome size for each accession was obtained using a CyFlow space cytometer (Partec Inc.). This instrument was equipped with a 488nm blue laser 50mW and a band-pass filter LP590 used as an emission filter. Also, prior to running the samples, gain and linearity of the instrument were adjusted by using DNA control PI from Sysmex. Finally, G1 peaks in *B. rapa* accessions and *P. sativum* were collected for each sample to calculate nuclear DNA content (1C in pg) and haploid genome size (in Mbp). Statistical tests were performed using R software v.3.5.1 (R Core Team, 2018).

## RESULTS

### Structural analysis of two highly contiguous Brassica rapa genomes

Structural comparison of the *B. rapa* ‘Z1’ genome against both *B. rapa* ‘Chiifu’ genome and B. napus ‘Darmor Bzh’ A subgenome revealed six large sequences specific to *B. rapa* ‘Z1’ on A01, A05 (two regions), A06, A08 and A09 (Table 1, see also Figure 1). These regions were qualified as ‘specific’ because no orthologous match could be found in the assembled *Brassica* genomes tested in this study. These six regions with a mean length of 12,179,136 bp (median = 9,811,381 bp) were composed of a mean number of TEs, rDNA and genes of 4,244.5 (median = 2,878), 25.5 (median = 19) and 160.5 (median = 135.5), respectively (Table 1). A total of 963 genes were found in the regions specific to ‘Z1’. On these 963 genes, 439 genes were only detected in the ‘Z1’ genome without any homologous similarity in ‘Chiifu’ genome. GO enrichment was performed for the genes localized in the ‘Z1’ specific regions, however no significant pathway was revealed. Similarly, we were not able to significantly detect enrichment of duplicated genes that may be localized in these regions. Moreover, these six ‘Z1’ regions contained an important percentage of ‘N’ sequences (mean = 30.59%, median = 26.38%). The observed ‘N’ percentage in these regions was associated with the scaffolding of long reads due to optical mapping. Thus, we might lack genetic information but thanks to optical mapping we are confident on the size of these structural variants.

**Table 1:**
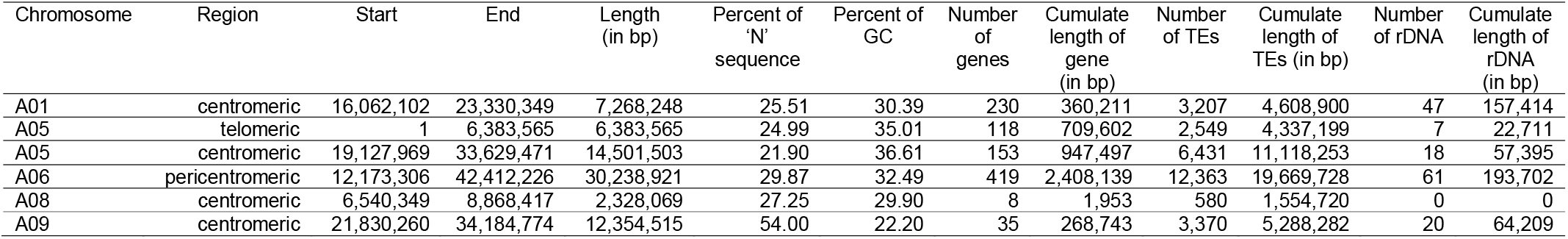
Position and length of the ‘Z1’ genome-specific regions (in comparison to *B. rapa* cv ‘Chiifu’). For each region, the number and cumulative length of genes, Transposable Elements and ribosomal DNA were identified.

**Figure 1:**
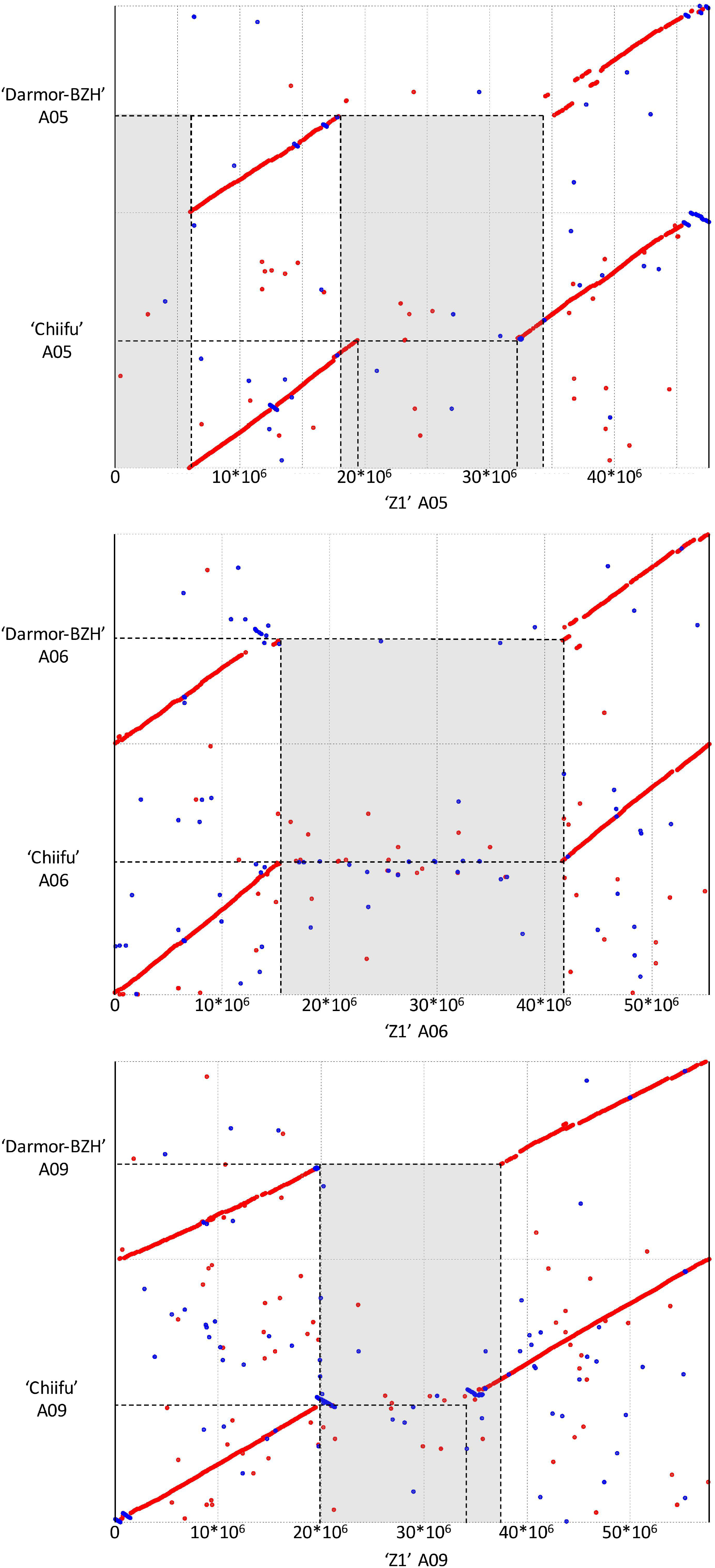
Dot plot comparing the assembly chromosome of *B. rapa* ‘Z1’ against *B. rapa* ‘Chiifu’ and *B. napus* Darmor Bzh A subgenome for the chromosomes A05 (A), A06 (B) and A09 (C). Grey boxes corresponded to regions specific to ‘Z1’ genome. Only alignment blocks with greater than 95% of identity and a length greater than 4,000 bp were shown.

Additionally, 81 inverted regions were identified on all chromosomes between ‘Z1’ (cumulative length = 23,066,463 bp) and ‘Chiifu’ (cumulative length = 176,703,831 bp) reciprocally (Supplementary Table 3). Inversions in ‘Z1’ and ‘Chiifu’ had a mean length of 281,771 bp (median = 72,650 bp) and 2,181,528 bp (median = 107,269 bp), respectively. A total of 22,274 (mean = 274.99, median = 14) and 21,780 (mean = 268.89, median = 13) genes were identified within ‘Z1’ and ‘Chiifu’ inversions, respectively. A total of 9,240 (mean = 114.07, median = 35) and 112,578 (mean = 1,389.85, median = 63) TEs were identified within ‘Z1’ and ‘Chiifu’ inversions, respectively. The cumulative length of TEs within the inversions corresponded to 49.86% and 44.15% of the total length of the inversions in ‘Z1’ and ‘Chiifu’, respectively. By contrast, a very low number of rDNA sequences were identified in these inversions (with overall 3 and 5 rDNA sequences in ‘Z1’ and ‘Chiifu’, respectively; Supplementary Table 3). Among these inversions, we were able to validate three inversions between ‘Z1’ and ‘Chiifu’, of which two were present on chromosome A05 (located from 6.48-6.50 Mb and 6.65-7.03 Mb) and one on chromosome A10 (located from 18.24-19.57 Mb) using PCR (Supplementary Figure 1).

### Repeat element content in B. rapa ‘Z1’ and ‘Chiifu’

The proportions of repeat elements (including Transposable Elements and ribosomal DNA) were estimated in *B. rapa* ‘Z1’ and *B. rapa* ‘Chiifu’. TEs were annotated along the ten chromosomes of both genomes with higher proportions in the centromeric regions. Considering a genome size of 529 Mb for *B. rapa* ‘Z1’ and ‘Chiifu’ after Johnston et al., (2005), TEs may represent 31.73% and 22.16% of ‘Z1’ and ‘Chiifu’ genomes respectively. However, considering the genome sizes determined in this study by flow cytometry (577 Mb and 545 Mb on average for ‘Z1’ and ‘Chiifu’, respectively, see below), TEs may represent 29.09% and 21.50% of ‘Z1’ and ‘Chiifu’ genomes respectively. LINEs and LTR (including Copia and Gypsy) TEs represent 3.02% and 8.58% of the ‘Z1’ genome, respectively. These proportions were also determined for the ‘Chiifu’ genome with 2.92% and 4.32% for LINEs and LTR TEs, respectively.

In *B. rapa* ‘Z1’, several chromosomes (especially the A05 and A06 chromosomes) had a higher proportion of TEs compared to ‘Chiifu’ assembled chromosomes. As expected, regions rich in TEs had a lower proportion of genes. Ribosomal sequences were observed in the ten chromosomes of both genomes with a higher proportion on the A01, A05, A06 and A09 compared to other chromosomes of ‘Z1’ (Figure 2). As the short arm of the chromosome A03 (bearing the Nucleolar Organizer Region, including a large domain of 45S rDNA) was not assembled, no rDNA were observed (Mun et al., 2010). These results were confirmed by CNV analysis, which allowed estimating the number of 45S rDNA copies to 1,416 and 1,750 in ‘Chiifu’ and ‘Z1’, respectively. Interestingly, the regions presenting a high number of rDNA and TEs corresponded to the six regions specific to the ‘Z1’ genome (Table 1, Figure 1). Proportions of each TE order in the two genomes were also investigated using χ^2^ test. No significant difference was observed between ‘Chiifu’ chromosomes, however, statistical differences were observed between ‘Z1’ chromosomes (α = 0.05). Indeed, chromosomes A05 and A06 had a higher percentage of LTR TEs in comparison to the other chromosomes (Figure 3, Supplementary Table 4). CNV analysis confirmed that the proportion of LTR Gypsy was higher in the ‘Z1’ genome (1,626 copies) compared to the ‘Chiifu’ genome (1,347 copies).

**Figure 2:**
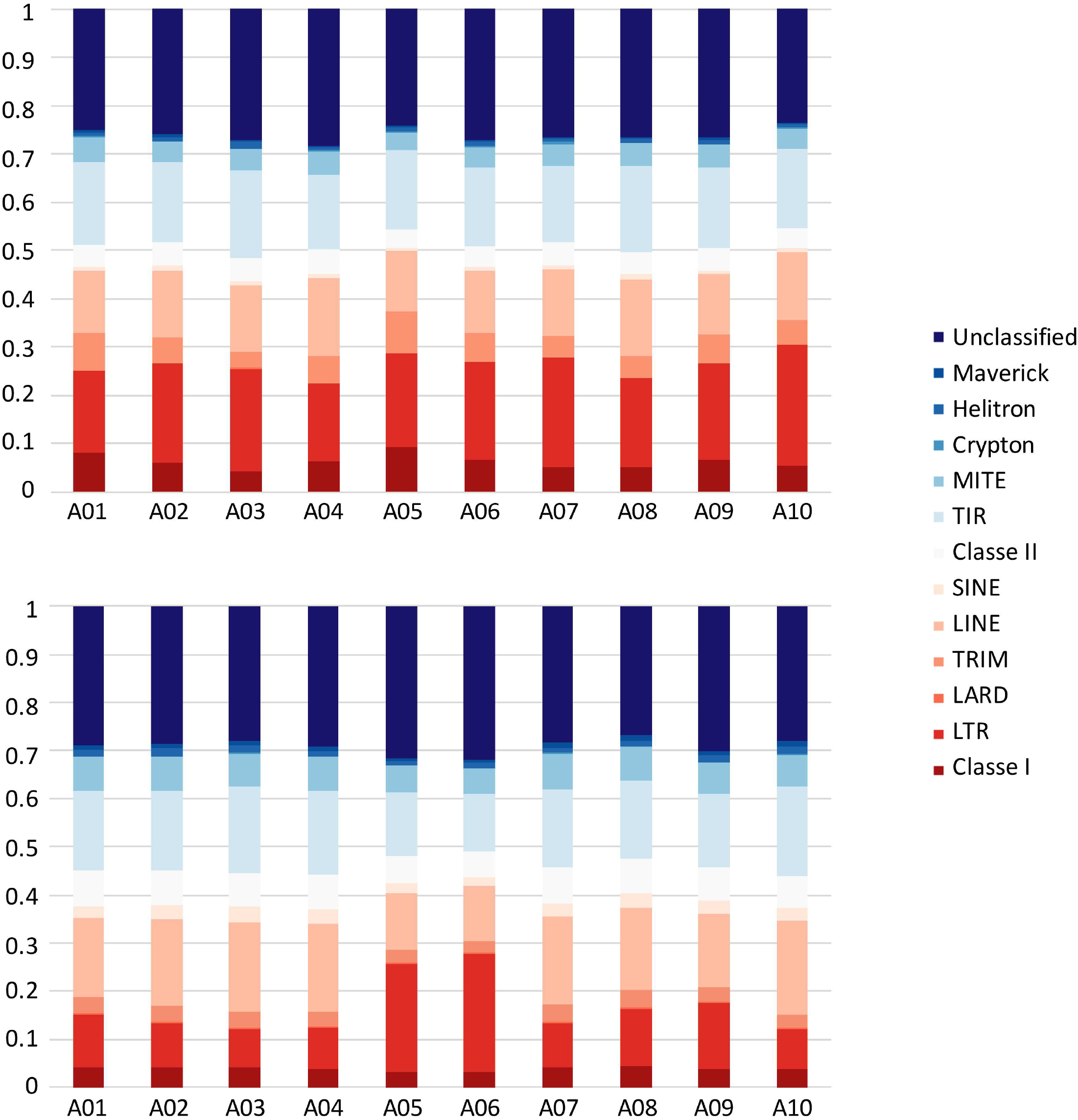
Circular diagrams of the chromosomes of (A) *B. rapa* ‘Chiifu’ and (B) *B. rapa* ‘Z1’. Black boxes represent chromosomes centromeric regions. Rings represent: (i) density of Transposable Elements, (ii) density of LTR, (iii) density of genes and (iv) presence-absence of ribosomal DNA (45S in blue and 5S in yellow).

**Figure 3:**
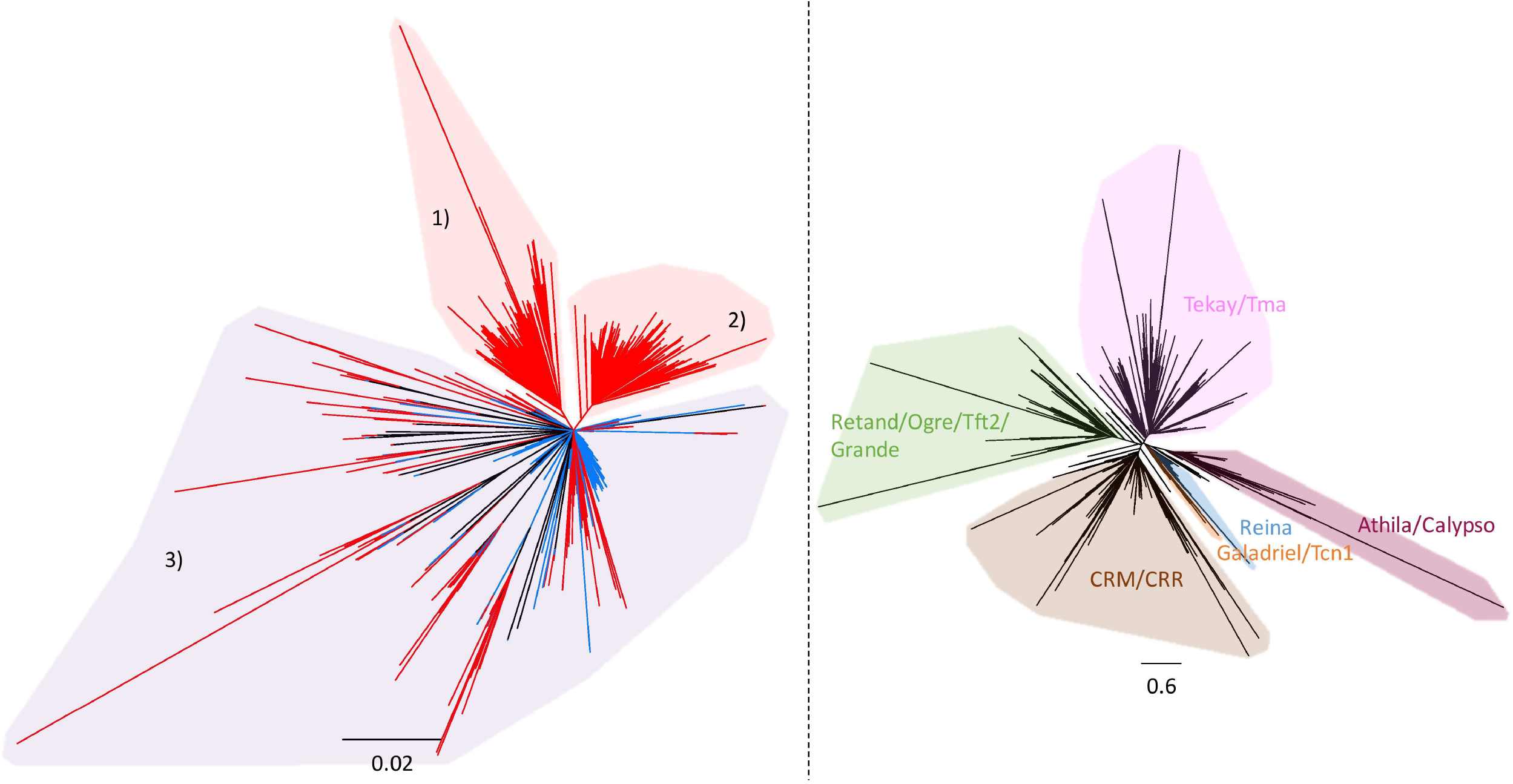
Transposable Element proportions in the 10 chromosomes of (A) *B. rapa* “Chiifu” and (B) *B. rapa* “Z1”. Class I and Class II corresponded to unclassified TEs within their respective Class.

As the large specific sequences of ‘Z1’ were made predominantly of rDNA and LTR Gypsy, phylogenies of both repeated elements were performed to explore their evolutionary dynamics. Ribosomal DNA phylogeny of the ‘Chiifu’ and ‘Z1’ genomes allowed us to identify three clades, one common to the two *B. rapa* samples (clade 3) and two containing only ‘Z1’ sequences (clades 1 and 2). These two clades contained 67 and 96 ‘Z1’ rDNA sequences, respectively, coming from chromosomes A01, A05, A06 and A09 (Figure 4, A). LTR Gypsy phylogeny allowed us to identify six major clades (Figure 4, B). Proportions of sequences within each LTR Gypsy clade was not significantly different between ‘Chiifu’ and ‘Z1’ (χ^2^ test; □ = 0.05).

**Figure 4:**
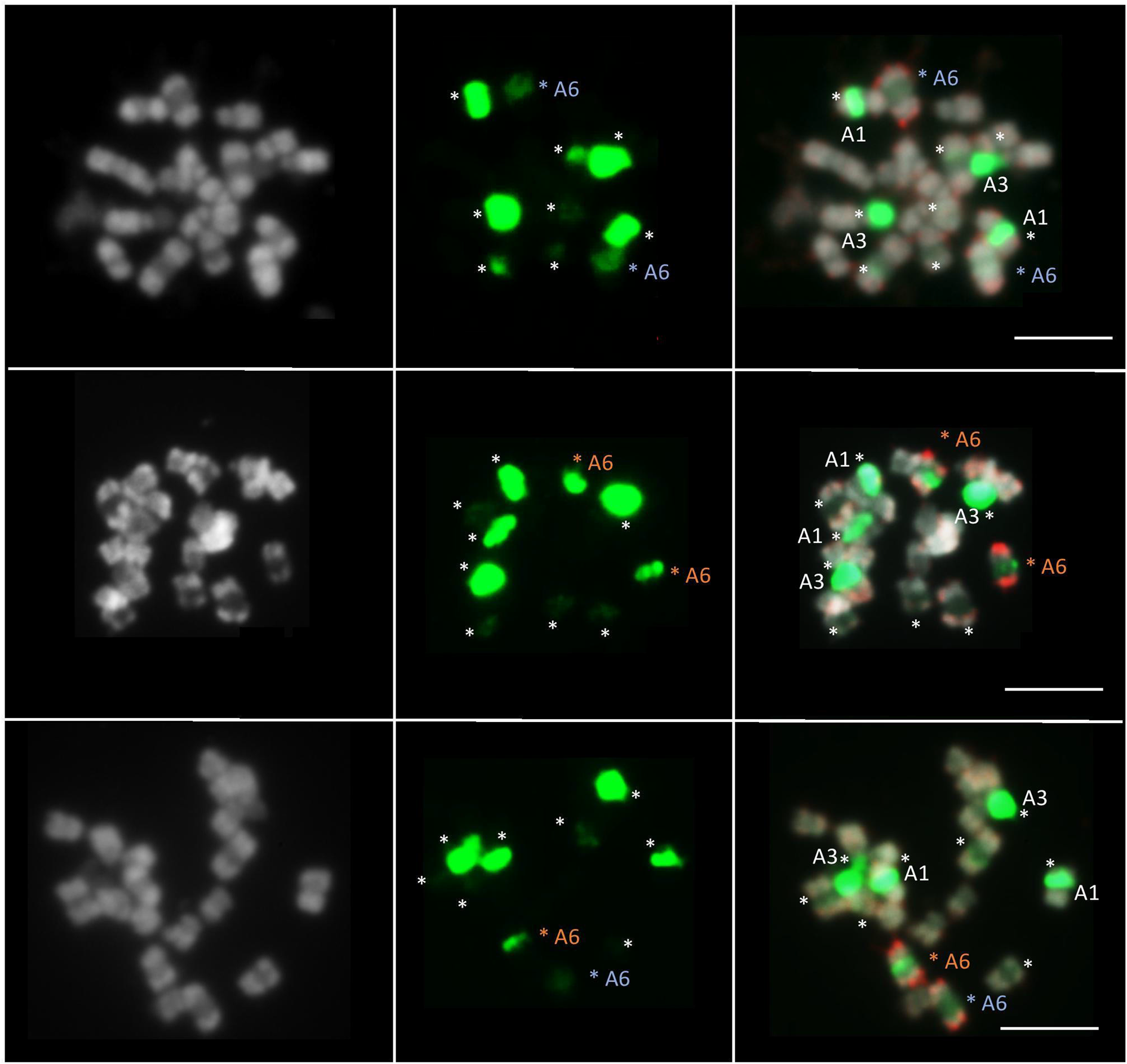
(A) Neighbour-joining phylogram of the rDNA of 168 *B. rapa* ‘Chiifu’ sequences (in blue) and 271 *B. rapa* ‘Z1’ sequences (in red). Clades 1 and 2 in red included only ‘Z1’ sequences. Clade 3 in purple, included ‘Chiifu’ and ‘Z1’ sequences. (B) Neighbour-joining phylogram and classification of LTR Gypsy elements of 2,447 *B. rapa* ‘Chiifu’ sequences, 2,908 *B. rapa* ‘Z1’ sequences and 60 Gypsy sequences from the Gypsy database for identifying major clades of TEs.

### Validation of the A06 chromosome length variation in B. rapa ‘Z1’ vs. ‘Chiifu’ using cytogenetic approaches

Fluorescence *in situ* hybridization (FISH) was performed on ‘Z1’, ‘Chiifu’ and F1 ‘Z1’x’Chiifu’ individuals to compare their A06 chromosomes, presenting high length variation. The 45S probe (green fluorescence) on *B. rapa* marked five different chromosome pairs (Figure 5). The strong FISH signal located on the A03 chromosomes likely reflected a large number of genes in the array of the 45S rDNA sequence. A second rDNA-rich locus was located on A01 chromosome, close to the centromere. BAC-FISH analysis with two specific BACs of A06 chromosome arms (red fluorescence) revealed that the 45S rDNA probe hybridized to a major site on *B. rapa* ‘Chiifu’ (Figure 5 A-C), a minor site on *B. rapa* ‘Z1’ (Figure 5D-F) and mix intensity sites, one major and one minor, on hybrid ‘Z1’x’Chiifu’ (Figure 5 G-I). Moreover, in *B. rapa*, the individual mitotic metaphase chromosome A06 ranged from 2.1 to 5.6 μm in length (observed from 14 metaphase sets). The chromosome A06 from ‘Chiifu’ was on average 17.6% (+/-6.96%) smaller compared to the chromosome A06 of ‘Z1’ (Supplementary Table 4).

**Figure 5:**
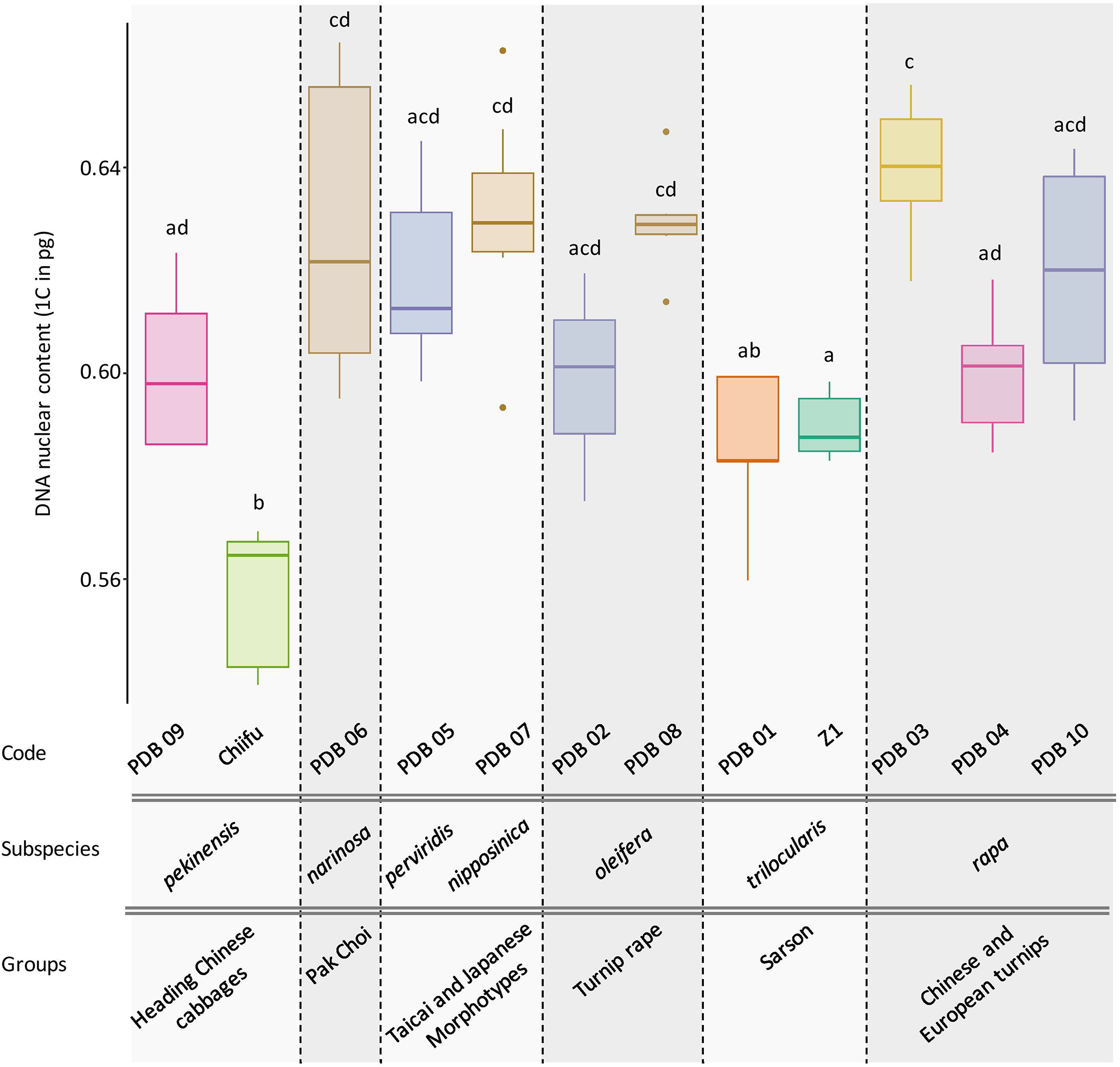
Fluorescent *in situ* hybridization was carried out using 45S rDNA (in green) and KBrB022P06 + KBrH003P24 BAC probes (A06) (in red). FISH analyses of somatic metaphase chromosomes was carried out in *B. rapa* ‘Z1’ (A-C), *B. rapa* ‘Chiifu’ (D-F) and Hybrid ‘Z1’x ‘Chiifu’ (G-I). The 45S rDNA probe on *B. rapa* marked five different chromosome pairs (stars). A06 chromosomes carrying 45S rDNA were indicated by a blue star in ‘Z1’ and an orange star in ‘Chiifu’. Chromosomes were counterstained with DAPI (grey). Bars represent 5 μm.

### Genome size variation and presence of inversions among B. rapa accessions

DNA content was obtained for ‘Z1’, ‘Chiifu’ and 10 accessions representing *B. rapa* core collection. After checking normality of data (Shapiro’s test, *P-value* = 0.487), the significative impact of accessions was modelised with an ANOVA including one factor (df = 11, *P-value* = 4.19e-13), followed by Tukey’s pairwise comparisons and group assignment based on *P-value* < 0.05. Of particular interest, the two focal species ‘Chiifu’ and ‘Z1’ have significantly different 1C DNA content with 0.557 (median = 0.565) and 0.590 (median = 0.588), respectively (Tukey HSD, *P-value* = 0.019). All accessions can be divided in seven subspecies and six groups structured within the core-collection (Figure 6). The variation in genome size does not seem to be associated with any of those phylogenetic structures.

**Figure 6:**
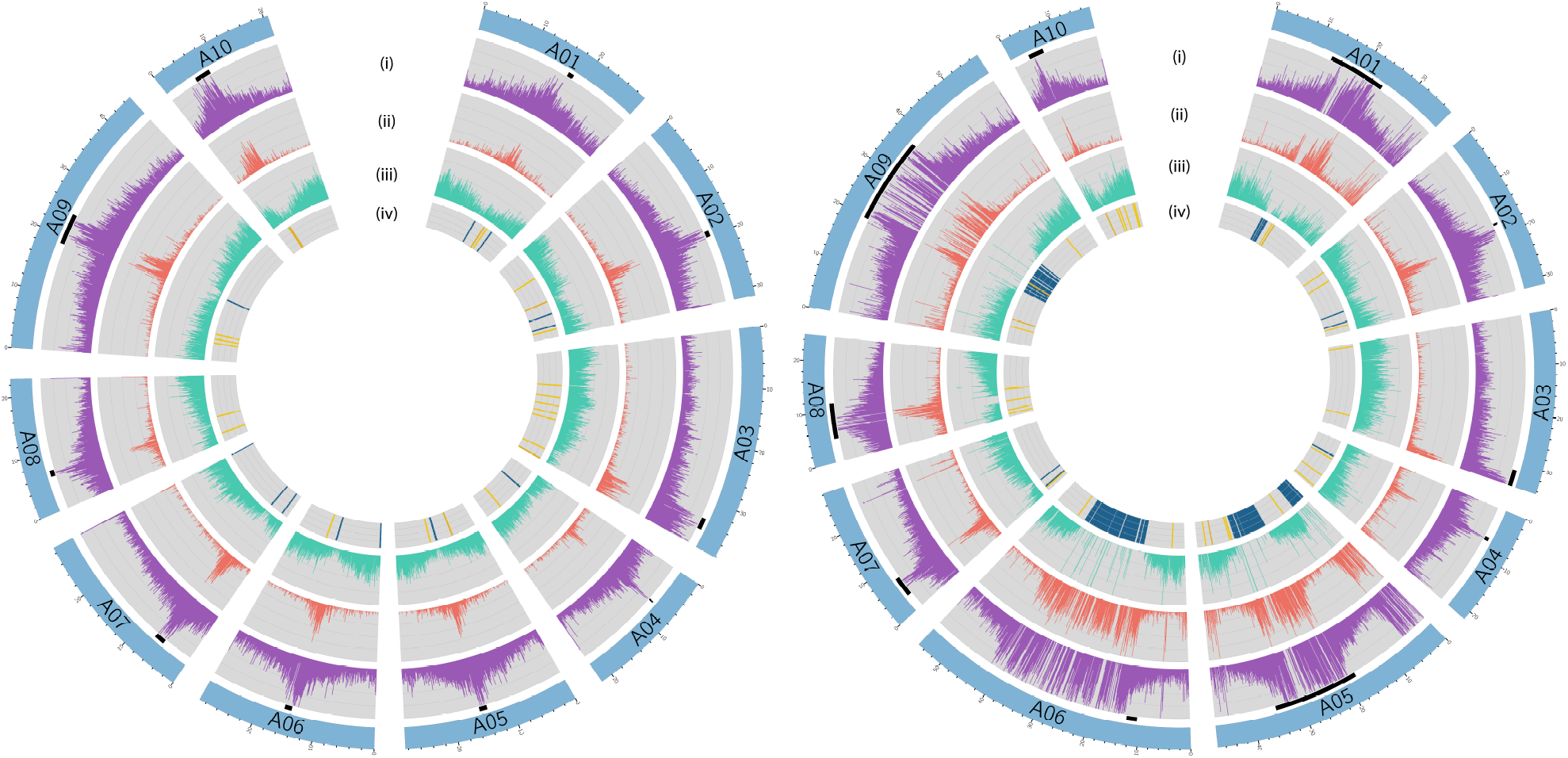
Box plot of the DNA nuclear content (1C) of 12 *B. rapa* accessions with 5 to 10 individuals per accessions. Colors indicate the Tukey’s pairwise comparisons and group assignment (□ = 0.05). Accessions are classified according to their higher order phylogeny (group and subspecies according to Cheng et al., 2016).

Finally, we explored the presence of these inversions in the same core-collection of *B. rapa* accessions. Although, it appeared that some nucleotide divergence might have prevented amplification, some accessions seem to share preponderantly either one or the other genetic structure (inversion on A05 zone II: populations 1, 2, 4, 5 and 10 might seem alike ‘Z1’ whereas populations 3, 7, 8 and 9 seem alike Chiifu; inversion on A10: populations 3, 4, 5, 6, 7, 9 and 10 seem more alike ‘Z1’ than).

## DISCUSSION

The large phenotypic diversity within *B. rapa* species has always triggered intensive investigations from the community. Ancient whole genome duplication followed by subgenome parallel selection has been shown to play a role in morphotype diversification in *B. rapa* (Cheng et al., 2016b). In addition, contrasted levels of cytosine methylations correlating with various TE densities have been proposed to shape subgenome dominance and fractionation levels within the genome of *B. rapa* (Cheng et al., 2016a). However, the extent of TE diversity within the species as well as the structural genomic variants that might additionally explain within species diversity have merely been touched upon.

### Detection of structural variations between two accessions of *B. rapa* and impact on genome size

A recent study on whole-genome resequencing of eight *B. napus* accessions revealed millions of genomic variants resulting in 6.8% to 13.2% (for a total of 77.2 Mb to 149.6 Mb) of the genome presenting such variations (Song et al., 2020). Similarly, chromosome-level assemblies of seven *A. thaliana* genomes unravel overall 13-17 Mb of rearranged genomic variants and up to 6.5 Mb (or 5.5% of the reference genome) of genome-specific sequence based on all possible pairwise comparisons (Jiao and Schneeberger, 2020). In the present study, we identified 23 Mb of inversions (representing 4.3% of the genome) and 73 Mb of specific insertions (representing 13.8% of the genome) in ‘Z1’ compared to ‘Chiifu’. These results show the large unexplored genomic diversity present in only two accessions of *B. rapa*. Although we cannot directly address their role in phenotypic diversification within the species, these insertions include 439 coding regions for which almost half of them cannot find an orthologous copy in the ‘Chiifu’ genome. These genes might be part of the dispensable gene pool as explained in a study gathering ten accessions of *B. oleracea* where 81.3% of genes represent the core whereas the remaining genes (18.7%) represent dispensable genes (Golicz et al., 2016).

Besides the functional consequences of such variants, the whole genome structure could be particularly impacted. The variation in genome sizes between our focal accessions has been previously addressed with estimations varying between 529 Mb and 443 Mb for ‘Z1’ and ‘Chiifu’ respectively (Belser et al., 2018; Zhang et al., 2018). However, the disparate methods used: either by flow cytometry or kmer analysis can reveal wide differences (see Zhang et al., 2018). Here, using internal control and repeated evaluations of DNA content by flow cytometry, we estimated an increase of ca. 6% of genome size between ‘Z1’ and ‘Chiifu’. In addition, we were able to validate the difference of the A06 chromosome lengths between the focal accessions using a ribosomal DNA probe in a BAC-FISH experiment showing first hand the possible additional impact of these elements on chromosome and genome size.

Extending our analysis to the *B. rapa* core-collection, we explored the variation in genome sizes among different accessions of *B. rapa* representatives of the phenotypic diversity in the species. Interestingly, up to 16% variation in genome size was observed among the 12 accessions without visible structuration according to their phylogenetic relationships. These differences are probably due to intraspecific variation in repeat content and insertion/deletion of sequences as their expansion correlates with the diversification of morphotypes (ca. 1mya, Zhao et al., 2013).

Very few studies tried so far to identify intra-specific genome size variation in Brassicaceae (with same ploidy level and same number of chromosomes). In Johnston et al., (2005), *B. rapa* species (including unknown accessions) exhibited wider standard error than any other Brassicaceae species. By comparison in *Arabidopsis thaliana*, the analyses of DNA content in ten ecotypes showed little variation with less than 1% among the different accessions (Johnston et al., 2005) which seemed to be corroborated with recent assemblies of various *A. thaliana* accessions with low genome size variance (Jiao and Schneeberger, 2020).

### Contrasted repeat landscapes between the focal accessions

Usually, genome size variation is often associated with TE content and ribosomal repeats as these elements could be highly dynamic. The large specific insertions identified in ‘Z1’ allowed to improve our understanding of centromeric and pericentromeric regions that are usually difficult to sequence and assemble in complex genomes such as plant genomes, due to their high density in repeats. Interestingly, apart from some dispensable genes, specific ‘Z1’ insertions exhibited mostly both rDNA copies and LTR elements.

In *Brassica* species, the number of 45S rDNA loci is variable. For example, *B. oleracea, B. nigra* and *B. rapa* have four, six and ten rDNA loci, respectively (Hasterok et al., 2001). However, the number of 45S (and 5S) rDNA loci varies among varieties within each species. Hasterok et al., (2006) identified ten 45S rDNA loci in all *B. rapa*, except in *B. rapa* subsp. *oleifera* (DC.) ‘Schneeball’ (6 loci). Here, we investigated the 45S rDNA in two other *B. rapa* varieties, ‘Z1’ and ‘Chiifu’. The number of 45S rDNA loci identified in these two varieties (10 loci, Figure 5) is identical to the number of 45S rDNA loci identified in the majority of other *B. rapa* varieties (Hasterok et al., 2006). However, based on the signal intensity, as shown by (Xiong and Pires, 2011) for the chromosome A05 of *B. rapa* ‘Chiifu’ vs. A05 of ‘IMB218’, the loci on the chromosome A06 in *B. rapa* ‘Z1’ had low signal intensity but showed a decondensation while the loci on the chromosome A06 in ‘Chiifu’ are fully condensed. As the A03 locus has been shown to be the only one carrying transcriptionally active ribosomal genes in the nucleolar organizer region (Hasterok and Maluszynska, 2000), it could be of interest to test if the 45S rDNA genes localized on the A06 in ‘Z1’ are transcribed and active.

Although the number of loci between ‘Z1’ and ‘Chiifu’ is identical, the number of 45S rDNA copies is different between these two varieties. The number of copies estimated in ‘Z1’ (1,750 copies) is highly similar to the number of rDNA copies previously estimated in this variety (1,709 copies, Sochorová et al., 2017) and is more abundant than the number of copies identified in ‘Chiifu’ (1,416 copies). This difference could be the result of rearrangements with loss of rDNA copies in *B. rapa* ‘Chiifu’, as demonstrated in the polyploid species such as *B. napus* or *Nicotiana tabacum* (Lim et al., 2000; Książczyk et al., 2011; Sochorová et al., 2017). However, rDNA phylogeny of ‘Chiifu’ and ‘Z1’ (Figure 4, A) has shown that among the three clades identified, two included only ‘Z1’ sequences (163 sequences). Thus, the most parsimonious scenario would indicate that these two clades most probably result from duplication events in ‘Z1’ after the divergence between the two focal accessions and not from loss of copies in ‘Chiifu’.

Regarding TE content, assembly of the ‘Z1’ genome has allowed a first comparative exploration of the repetitive compartment between ‘Z1’ and ‘Chiifu’ (Belser et al., 2018), however differences were mainly attributed to methodological dissimilarities between genome assemblies. Here, using a consistent analysis pipeline for TE detection and annotation in these two accessions, we can reveal the impact of TEs on *B. rapa* intraspecific diversity taking into account the structural variations.

Using genome size estimation for *B. rapa*, (i.e 529Mb according to Johnston et al., 2005), TEs represent 31.73% and 22.16% of ‘Z1’ and ‘Chiifu’ genomes, respectively. However, considering the genome sizes obtained in this study (577 Mb and 545 Mb on average for ‘Z1’ and ‘Chiifu’, respectively), TEs may represent 29.09% and 21.50% of the genome in ‘Z1’ and ‘Chiifu’, respectively. Overall, our methodology allowed us to retrieve similar numbers of TEs compared to what is expected with 29% and 24% of TE in accessions ‘Z1’ and ‘Chiifu’, respectively (Belser et al., 2018). Interestingly, a tangible difference in TE content is consistently observed between these two accessions and we proposed that this pattern is not widely distributed along the genome but focused on specific chromosomes and identified structural variants.

The detailed analysis of the structural variants in ‘Z1’ showed that while inversions do not significantly sustain a higher proportion of TEs compared to the rest of the genome, specific insertions in ‘Z1’ carry a large proportion of LTR Gypsy and LINEs. TE proportions could be underestimated in these regions as a large part of the inserted sequences were unsequenced data, we are nonetheless confident on the length of these sequences thanks to reliable optical mapping scaffolds.

TE intraspecific diversity has been previously shown to be driven by an expansion of recent LTR retrotransposons in *B. rapa* with rapid nucleotide substitution (Zhao et al., 2013). However, we do not observe such patterns using a phylogenetic approach. Instead, all LTR Gypsy clades seem to be represented in the inserted regions. This tendency seems to be particularly exacerbated in *B. rapa* and by extension in the A subgenome of *B. napus* compared to *B. oleracea* and the C subgenome of *B. napus* as demonstrated in Song et al., (2020). In addition, the inserted LTR retrotransposons do occur in clusters on chromosomes A05, A06 and A09 and their spreading predates the divergence between *B. rapa* and *B. oleracea* (Song et al., 2020).

## CONCLUSION and PERSPECTIVES

Our results unveil the necessity for long-read sequencing strategies and developing various whole-genome references for agronomically important species (Danilevicz et al., 2020). This recent endeavour in constructing pangenomes is revealing the importance of structural variants and the diversity in repeat content within species. This is especially relevant in species showing broad phenotypic diversity such as *Brassica sp*. Besides the role of these variants in *Brassica* diversification we can only wonder at the role of such variants in other processes such as recombination. In that case, structural rearrangements (especially inversions) act as modifiers of recombination that create different patterns of recombination rate across the genome (Ortiz-Barrientos et al., 2016) leading to segregation distortion and partial lethality (Lynch and Force, 2000; Fransz et al., 2016). Similarly, TEs have been shown to cause post-zygotic barriers and reproductive isolation in a diversity of taxa (reviewed in Serrato-Capuchina and Matute, 2018). Thus, continuing the investigation of the diversity within *B. rapa* is an important avenue to address. First hints suggest that both structural variants and repeat content seem highly dynamic among the different accessions of *B. rapa* composing the core-collection. Finally, exploring this diversity will deeply improve our understanding of the species adaptive processes and could be used in the creation of new hybrid varieties.

## Author Contributions

JFC and JB conceived and designed the experiments with inputs from LM. TC, SL and LM implemented the REPET pipeline and performed the work on TE annotation. JB, LM, CF and JM performed the bioinformatic analyses. J-MA and CB analysed the generated optical maps and verified regions of interests. AB, VH and OC performed the cytological analyses. GT performed the flow-cytometry experiments. GD and ML performed the DNA extractions and PCR to verify inversions. A-MC and MR-G helped supervise the project and participated in critical thinking of the results with LM, FB, CF, OC, JB and JFC. JFC and JB wrote the manuscript; all authors approved the final manuscript.

## Acknowledgements

This work was made possible by financial support of the European Union and a Marie Sklodowska-Curie grant (number: 791908) awarded to JFC. This work was additionally supported by the France Génomique National infrastructure, funded as part of the « Investissements d’Avenir » program managed by the Agence Nationale pour la Recherche (contract ANR-10-INBS-09). We acknowledge the BrACySol BRC (INRA Ploudaniel, France) that provided us with the seeds of *B. rapa* accessions and Marie-Madeleine Gilet for growing and taking care of the accessions. We would also like to thank all the technical staff of the greenhouse for management of the plant material (especially L. Charlon, P. Rolland, J.-P. Constantin, J.-M. Lucas and F. Letertre).

## Contribution to the Field Statement (to be included in online submission 200 words)

Using comparative genomics and cytogenetics approaches, we explored the structural consequences of the presence of large repeated sequences in two genomes of *B. rapa*. These two genomes representing divergent accessions referred to ‘Z1’ and ‘Chiifu’ exhibited inversions, insertions as well as differences in TE and rDNA copy numbers largely influencing chromosome size and global DNA content. In addition, we assessed presence/absence inversions and diversity in genome size among additional accessions of *B. rapa* representing the whole range of morphotypes within the species. We hypothesized that these structural differences could have participated in *B. rapa* phenotypic diversity and may play a role in preventing recombination in these regions.

## SUPPLEMENTARY TABLES

Supplementary Table 1: Accession identification and origin of the *B. rapa* accessions.

Supplementary Table 2: Primer identifiers, sequences and target amplification.

Supplementary Table 3: Position and length of the *B. rapa* ‘Chiifu’ vs ‘Z1’ inversions. For each region, the number and cumulative length of genes, Transposable Elements and ribosomal DNA were identified.

Supplementary Table 4: Transposable Element proportions in the 10 chromosomes of *B. rapa* ‘Chiifu’ and *B. rapa* ‘Z1’. Class I and Class II correspond to TEs for which order’s annotation remains unknown.

Supplementary Table 5: A06 individual chromosome lengths (in μm) at mitotic metaphase in F1 hybrid (‘Z1’x‘Chiifu’).

## SUPPLEMENTARY FIGURE

Supplementary Figure 1: PCR amplifications and agarose gels validating inversions in three genomic regions on (A) chromosome A05 first inversion, (B) chromosome A05 second inversion and (C) chromosome A10. Both sides of each inversion have been investigated (L for left side and R for right side) in the focal accessions and *B. rapa* core collections (named here as PBD1 to PBD10), ordered on the gel as follows: ‘Chiifu’, ‘Z1’, PBD001 to PBD010.

