## Supplementary figures and images for "Large genomic variants reveal unexplored intraspecific diversity in *Brassica rapa* genomes"

### Supplemental Figure 1

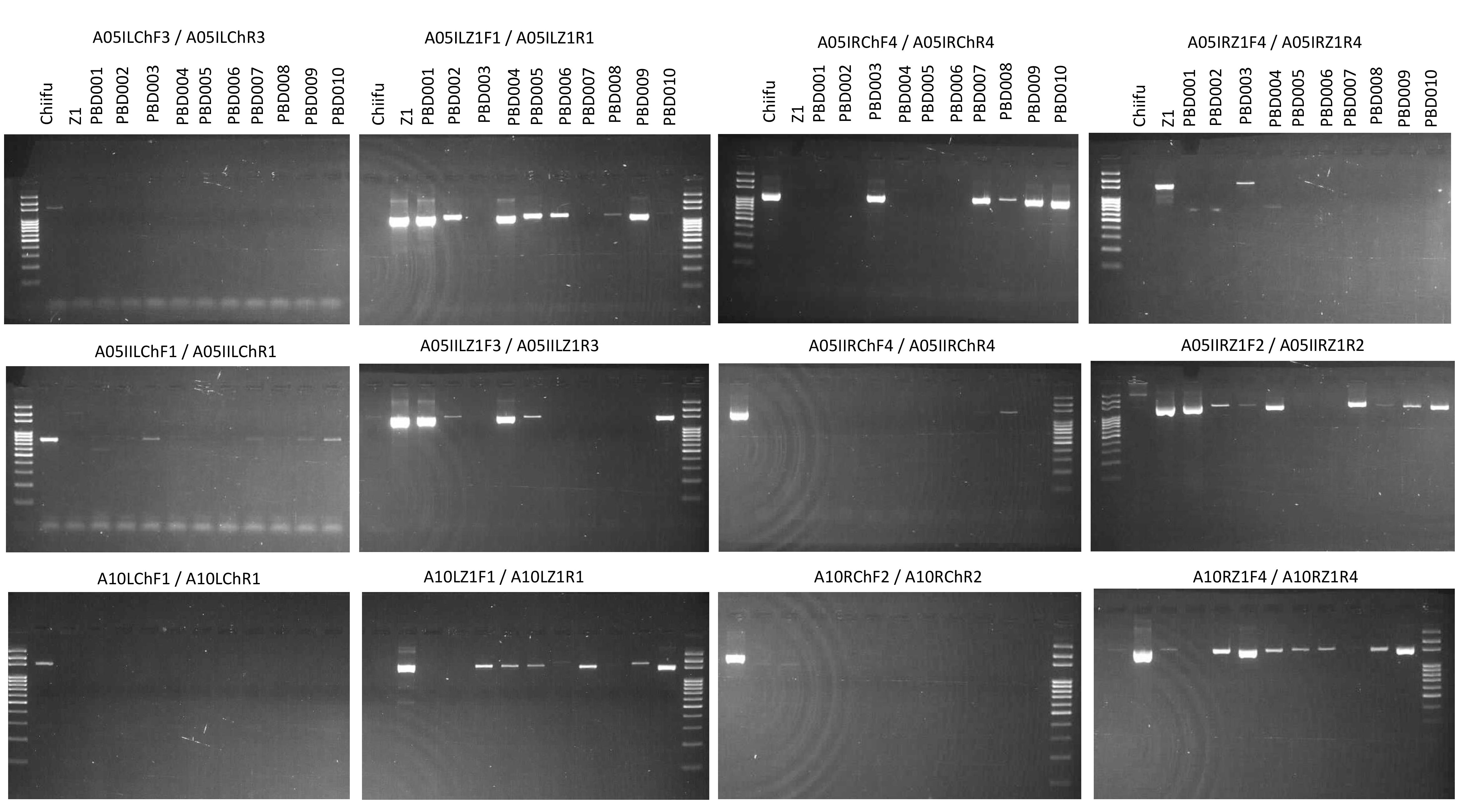
