## Supplemental Table 4 for "Large genomic variants reveal unexplored intraspecific diversity in *Brassica rapa* genomes"

Supplementary Table 4: Transposable Element proportions in the 10 chromosomes of *B. rapa* ‘Chiifu‘ and *B. rapa* ‘Z1‘. Class I and Class II correspond to TEs for which order’s annotation remains unknown.

|  | Class I | LTR | | LARD | TRIM | DIRS | Penelope | LINE | SINE | Class II | TIR | MITE | Crypton | Helitron | Maverick | Unclassified |
| --- | --- | --- | --- | --- | --- | --- | --- | --- | --- | --- | --- | --- | --- | --- | --- | --- |
| *B. rapa* “Chiifu” | | |  |  |  |  |  |  |  |  |  |  |  |  |  |  |
| A01 | 0,0825 | 0,1698 | | 0,0005 | 0,0771 | 0,0000 | 0,0000 | 0,1293 | 0,0074 | 0,0458 | 0,1711 | 0,0527 | 0,0010 | 0,0080 | 0,0055 | 0,2493 |
| A02 | 0,0606 | 0,2053 | | 0,0009 | 0,0530 | 0,0000 | 0,0000 | 0,1392 | 0,0097 | 0,0503 | 0,1660 | 0,0426 | 0,0004 | 0,0089 | 0,0038 | 0,2592 |
| A03 | 0,0420 | 0,2142 | | 0,0003 | 0,0356 | 0,0000 | 0,0000 | 0,1363 | 0,0097 | 0,0463 | 0,1818 | 0,0455 | 0,0010 | 0,0129 | 0,0041 | 0,2702 |
| A04 | 0,0646 | 0,1593 | | 0,0006 | 0,0582 | 0,0000 | 0,0000 | 0,1589 | 0,0108 | 0,0495 | 0,1561 | 0,0489 | 0,0005 | 0,0068 | 0,0041 | 0,2816 |
| A05 | 0,0931 | 0,1939 | | 0,0015 | 0,0865 | 0,0000 | 0,0000 | 0,1250 | 0,0068 | 0,0374 | 0,1646 | 0,0369 | 0,0009 | 0,0104 | 0,0034 | 0,2395 |
| A06 | 0,0672 | 0,2011 | | 0,0006 | 0,0603 | 0,0000 | 0,0000 | 0,1292 | 0,0081 | 0,0426 | 0,1643 | 0,0422 | 0,0008 | 0,0100 | 0,0037 | 0,2701 |
| A07 | 0,0508 | 0,2285 | | 0,0002 | 0,0439 | 0,0000 | 0,0000 | 0,1384 | 0,0093 | 0,0461 | 0,1574 | 0,0456 | 0,0051 | 0,0061 | 0,0042 | 0,2644 |
| A08 | 0,0515 | 0,1849 | | 0,0013 | 0,0446 | 0,0000 | 0,0000 | 0,1591 | 0,0103 | 0,0462 | 0,1789 | 0,0456 | 0,0002 | 0,0087 | 0,0037 | 0,2650 |
| A09 | 0,0652 | 0,2020 | | 0,0008 | 0,0586 | 0,0000 | 0,0000 | 0,1238 | 0,0088 | 0,0470 | 0,1676 | 0,0466 | 0,0000 | 0,0087 | 0,0051 | 0,2657 |
| A10 | 0,0559 | 0,2491 | | 0,0013 | 0,0493 | 0,0000 | 0,0000 | 0,1416 | 0,0072 | 0,0429 | 0,1655 | 0,0423 | 0,0007 | 0,0063 | 0,0026 | 0,2354 |
| *B. rapa* “Z1” | |  | |  |  |  |  |  |  |  |  |  |  |  |  |  |
| A01 | 0,0405 | 0,1121 | | 0,0021 | 0,0338 | 0,0000 | 0,0000 | 0,1631 | 0,0259 | 0,0730 | 0,1656 | 0,0710 | 0,0008 | 0,0139 | 0,0098 | 0,2883 |
| A02 | 0,0427 | 0,0910 | | 0,0024 | 0,0349 | 0,0000 | 0,0000 | 0,1778 | 0,0305 | 0,0719 | 0,1644 | 0,0718 | 0,0008 | 0,0167 | 0,0105 | 0,2847 |
| A03 | 0,0400 | 0,0828 | | 0,0024 | 0,0325 | 0,0000 | 0,0000 | 0,1858 | 0,0321 | 0,0694 | 0,1808 | 0,0686 | 0,0009 | 0,0160 | 0,0104 | 0,2783 |
| A04 | 0,0373 | 0,0869 | | 0,0026 | 0,0304 | 0,0000 | 0,0000 | 0,1832 | 0,0308 | 0,0707 | 0,1752 | 0,0696 | 0,0009 | 0,0133 | 0,0092 | 0,2899 |
| A05 | 0,0335 | 0,2246 | | 0,0018 | 0,0273 | 0,0000 | 0,0000 | 0,1175 | 0,0191 | 0,0566 | 0,1325 | 0,0558 | 0,0003 | 0,0095 | 0,0056 | 0,3159 |
| A06 | 0,0317 | 0,2468 | | 0,0019 | 0,0258 | 0,0000 | 0,0000 | 0,1125 | 0,0191 | 0,0532 | 0,1200 | 0,0526 | 0,0006 | 0,0101 | 0,0065 | 0,3192 |
| A07 | 0,0418 | 0,0937 | | 0,0020 | 0,0347 | 0,0000 | 0,0000 | 0,1837 | 0,0279 | 0,0740 | 0,1616 | 0,0731 | 0,0030 | 0,0104 | 0,0107 | 0,2834 |
| A08 | 0,0453 | 0,1178 | | 0,0031 | 0,0364 | 0,0000 | 0,0000 | 0,1723 | 0,0296 | 0,0720 | 0,1599 | 0,0712 | 0,0011 | 0,0124 | 0,0108 | 0,2681 |
| A09 | 0,0378 | 0,1380 | | 0,0020 | 0,0314 | 0,0000 | 0,0000 | 0,1522 | 0,0271 | 0,0687 | 0,1516 | 0,0677 | 0,0003 | 0,0137 | 0,0091 | 0,3003 |
| A10 | 0,0384 | 0,0827 | | 0,0026 | 0,0296 | 0,0000 | 0,0000 | 0,1944 | 0,0255 | 0,0673 | 0,1840 | 0,0670 | 0,0007 | 0,0171 | 0,0115 | 0,2792 |
