## Supplemental Table 5 for "Large genomic variants reveal unexplored intraspecific diversity in *Brassica rapa* genomes"

|  | Set 1 | Set 2 | Set 3 | Set 4 | Set 5 | Set 6 | Set 7 | Set 8 | Set 9 | Set 10 | Set 11 | Set 12 |
| --- | --- | --- | --- | --- | --- | --- | --- | --- | --- | --- | --- | --- |
| ‘Z1’ | 3.455 | 2.779 | 3.246 | 2.807 | 2.432 | 3.157 | 3.321 | 2.555 | 2.838 | 2.691 | 2.153 | 3.069 |
| ‘Chiifu’ | 2.484 | 2.589 | 2.383 | 2.422 | 2.134 | 2.674 | 2.471 | 2.04 | 2.327 | 2.43 | 1.919 | 2.493 |

Supplementary Table 4: A06 individual chromosome lengths (in µm) at mitotic metaphase in F1 hybrid (‘Z1‘*‘Chiifu’).
